# hsa-miR-130b-3p Predicts Poor Prognosis Across Renal Cell Carcinoma Subtypes via CD8⁺T Cell Depletion: A Multi-Omics and Clinical Validation StudyImmune Evasion

**DOI:** 10.1101/2025.09.15.676345

**Authors:** Weilong Yang, Zhong Dong

## Abstract

**Abstract Background:** Renal cell carcinoma (RCC) is an aggressive genitourinary malignancy comprising three major subtypes: clear cell (KIRC), chromophobe (KICH), and papillary (KIRP). MicroRNAs (miRNAs) orchestrate renal cell carcinoma (RCC) progression, yet pan-subtype biomarkers remain elusive.

**Methods:** Hsa-miR-130b-3p was upregulated in all subtypes (log₂FC=1.65,FDR=1.07×10⁻⁵ in GSE16441), correlating with poor OS (HR=1.56-2.38, p<0.05) and reduced CD8+ T cell infiltration (r=-0.32, p=0.01). Functional enrichment linked it to mTOR/HIF-1 signaling.

**Results:** Hsa-miR-130b-3p was upregulated in all subtypes (log₂FC=1.65, FDR=1.07×10⁻⁵ in GSE16441), correlating with poor OS (HR=1.56-2.38, p<0.05) and reduced CD8+ T cell infiltration (r=-0.32, p=0.01). Functional enrichment linked it to mTOR/HIF-1 signaling.

**Conclusion:** hsa-miR-130b-3p emerges as a simple yet powerful predictor of RCC progression and prognosis, warranting further clinical validation. As a pan-subtype biomarker (3-year AUC=0.82), hsa-miR-130b-3p may facilitate RCC immune evasion, warranting liquid biopsy validation.

This study fills a gap in non-clear cell RCC (KICH/KIRP) by identifying hsa-miR-130b-3p as the first pan-subtype miRNA biomarker (3-year AUC=0.82, p<0.001), validated in 894 TCGA samples and 34 GSE16441 clinical cases. (See **Figure 1** for the Graphical Abstract.)

Figure 1
Graphical Abstract

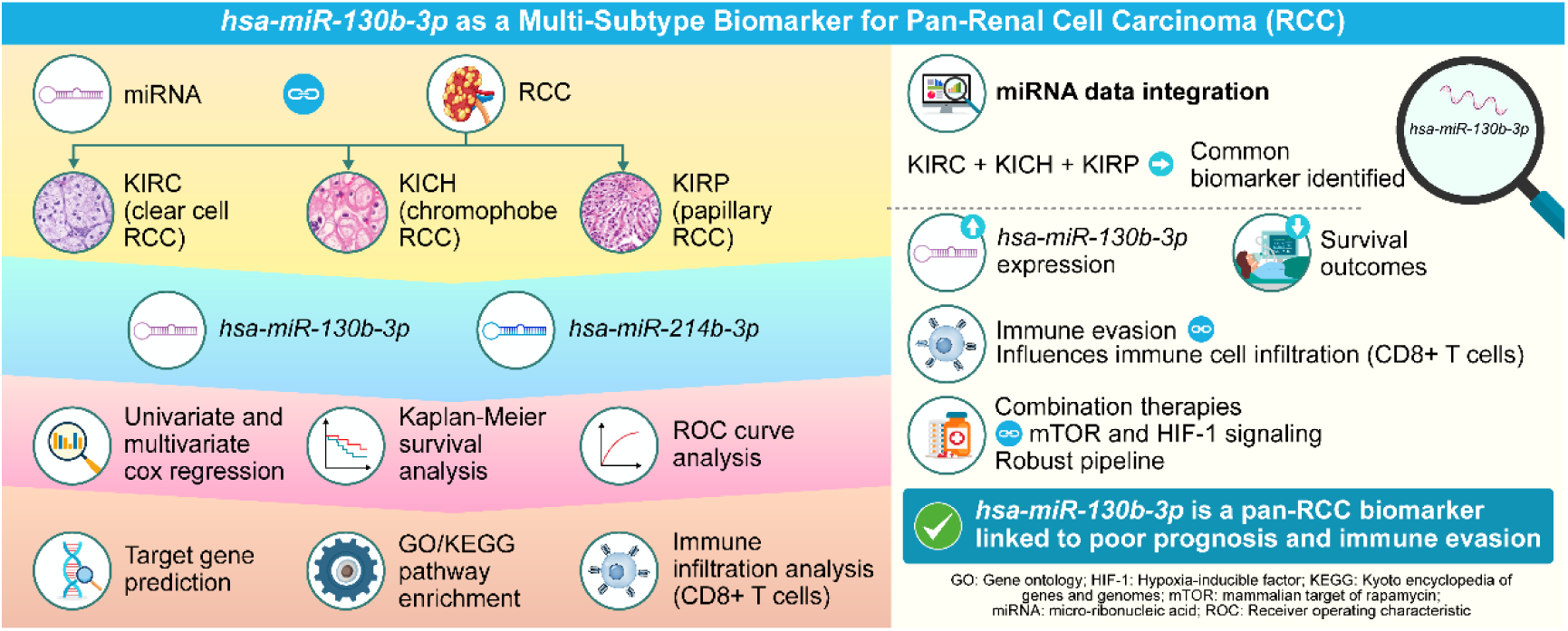

## Introduction

MicroRNA-mediated post-transcriptional regulation is now recognized as a key accelerator of renal cancer spread. For instance, elevated miR-34a fosters RCC metastasis via MET pathway activation.^2^ Nevertheless, prior investigations have largely concentrated on individual RCC subtypes, neglecting conserved miRNA networks across subtypes and their interplay with the immune microenvironment. Consequently, identifying miRNA signatures conserved among KIRC, KICH, and KIRP is imperative for refining precision diagnostics and therapeutics.^11^

Recently, CIMB’s special issue Non-coding RNA in Cancer highlighted the need for pan-subtype miRNA biomarkers in RCC (DOI:10.3390/cimb47030045). Here, we identify hsa-miR-130b-3p as a novel pan-subtype regulator linking miRNA dysregulation to CD8+ T cell immunosuppression, addressing this unmet need.

## Materials and methods Data acquisition

Transcript-level miRNA counts (TMM-normalized) for TCGA-KIRC, TCGA-KICH and TCGA-KIRP were retrieved from the CancerMIRNome portal,^2^ together with matched mRNA and clinical follow-up information. To corroborate miRNA– mRNA interactions, we additionally downloaded level-3 DNA gene-expression matrices (HTSeq-Counts) for the same three TCGA cohorts from the GDC Data Portal. An external validation set was provided by the GSE16441 series, which contains paired miRNA profiles from 17 clear-cell RCC tumors and 17 adjacent normal renal cortex samples.

1. TCGA data (KIRC, KICH, KIRP transcriptomic and clinical data) and UCSC data were accessed in November 2024; GEO data (GSE16441) were accessed in June 2025.
2. No. All data retrieved from TCGA and GEO are fully de-identified, with no direct or indirect personal identifiers available. The authors did not and could not access any information that would allow identification of individual participants.

Integrating these multi-layer datasets enabled a comprehensive interrogation of hsa-miR-130b-3p, its putative targets and immune-infiltration signatures across the major RCC subtypes.

## Analysis of differential expressions

Differential miRNAs were pinpointed via edgeR v4.4.1 through tumor-versus-normal comparison.

## Univariate Cox regression

Univariate Cox models were employed to single out miRNAs associated with survival endpoints.

## Intersection analysis

Overlapping miRNAs in each RCC subtype were identified for differential expression and Cox regression analyses, and cross-subtype intersections were performed to identify miRNAs that might be targets for multi-subtype RCC.

## Multivariate Cox regression

Multivariate Cox analyses adjusted for age, gender, TNM stage, and tumor grade were performed using R (v4.4.1) to validate the independent prognostic values of the candidate miRNAs.

## Independent validation cohort (GEO2R, GSE16441)

To check whether hsa-miR-130b-3p holds up outside TCGA, we downloaded the GSE16441 set (Agilent GPL8659, 34 paired samples: 17 ccRCC tumours and 17 matched normal renal cortex). The workflow was straightforward: Import the raw intensity matrix with GEO query 2.66.0.Apply normexp background correction, quantile normalization and log₂ transformation.Run a paired moderated t-test (limma-voom) with Benjamini–Hochberg FDR ≤ 0.05 to compare tumour versus normal.

Pull the probe A_25_P00010437 (hsa-miR-130b-3p) and record its log₂FC and FDR.The result was clear: hsa-miR-130b-3p was markedly higher in tumours (log₂FC = 1.64802, FDR = 1.07 × 10⁻⁵).

## Kaplan-Meier analysis

Kaplan–Meier curves were plotted with survival using Sangerbox; significance was gauged by log-rank tests.

## ROC analysis

ROC curves were constructed using Sangerbox to benchmark diagnostic performance of the target miRNAs in combination with the three RCC subtypes.

## Target prediction and functional enrichment

The target genes of the miRNAs were predicted using TargetScan and analyzed for GO/KEGG enrichment using Sangerbox.

## Immune infiltration analysis

The type and infiltration extent of target miRNAs in immune cells in three RCC subtypes were assessed using the GSVA and CIBERSORT algorithms in R v4.4.1.

## Results

### Differential expression and prognostic screening

Sequencing data normalized to the TMM of TCGA miRNAs for the three RCC subtypes (TCGA-KIRC, TCGA-KICH, and TCGA-KIRP) and clinical survival of the relevant samples were first obtained from the CancerMIRNome website and preprocessed.

The miRNA sequencing expression data of the three RCC subtypes were analyzed using the edgeR package in R (v4.4.1), and miRNAs differentially expressed between the tumor and normal tissue groups (DEmiRs) were identified. The differentially expressed miRNAs with logFC >0.5 and p < 0.05 were considered upregulated.

One-way Cox regression analysis of the miRNA expression data from the three RCC subtypes and the clinical survival information of the samples was performed to screen prognostic miRNAs with two outcomes: OS and PFI. HR >1 and p <0.05 were the screening conditions. The identified miRNAs are potential prognostic biomarkers for each RCC subtype and outcome.^12^(See **Figure 2** for the heatmap results of differential expression analysis, **Figure 3** for the volcano plot results of differential expression analysis, and **Figure 4** for the forest plot results of univariate Cox analysis.)

**Figure 2.**
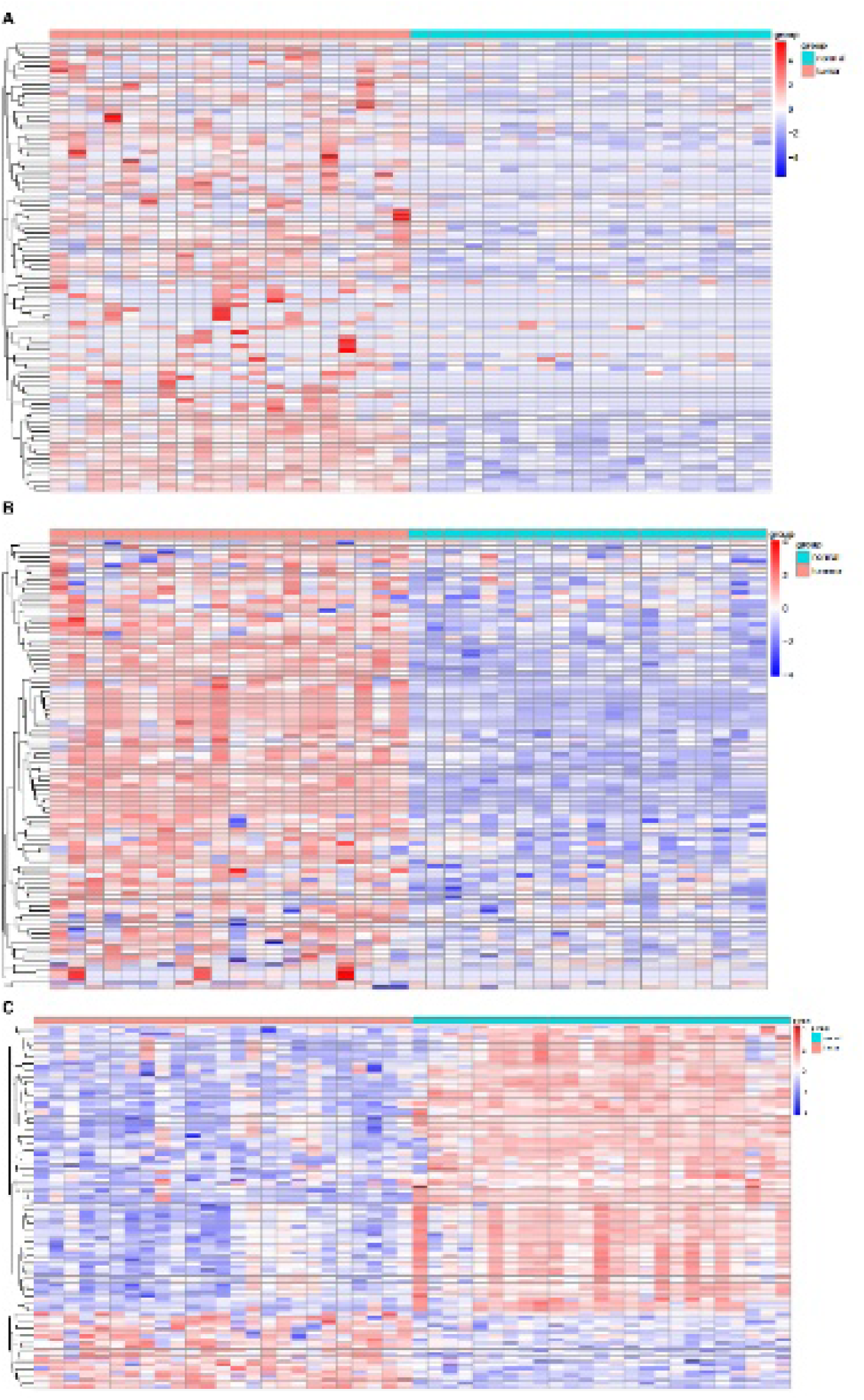
(A-C) Differential expression heatmaps of miRNAs in KIRC, KICH, and KIRP (tumor vs. normal). Analyses were performed using edgeR v4.4.1 (logFC>0.5, p<0.05 for upregulated miRNAs). Color gradients represent relative miRNA expression levels (red=high, blue=low); clustering identifies subtype-specific and shared differential miRNAs. hsa-miR-130b-3p (marked in red boxes) was consistently upregulated across all subtypes, laying the foundation for subsequent pan-subtype analysis (Figure 5).

**Figure 3.**
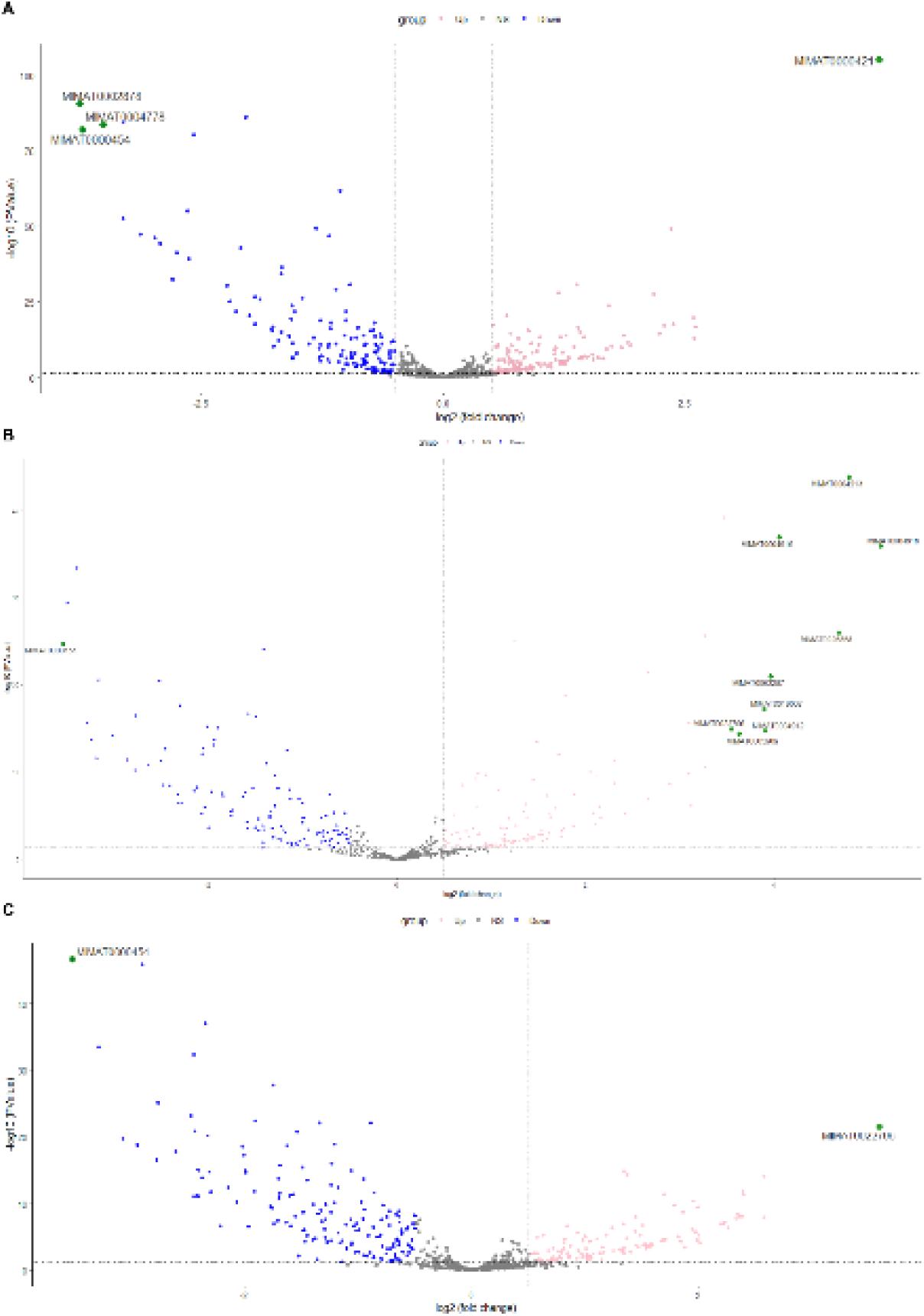
(A-C) Volcano plots of differentially expressed miRNAs in KIRC, KICH, and KIRP (tumor vs. normal). X-axis=log₂(fold change), Y-axis=-log₁₀(Padj); red=upregulated (Padj<0.05), blue=downregulated (Padj<0.05), gray=non-significant. Arrows highlight hsa-miR-130b-3p, which showed consistent upregulation (log₂FC≈1.65, Padj<0.001) across subtypes—consistent with its validation in GSE16441 (Figure 9).

**Figure 4.**
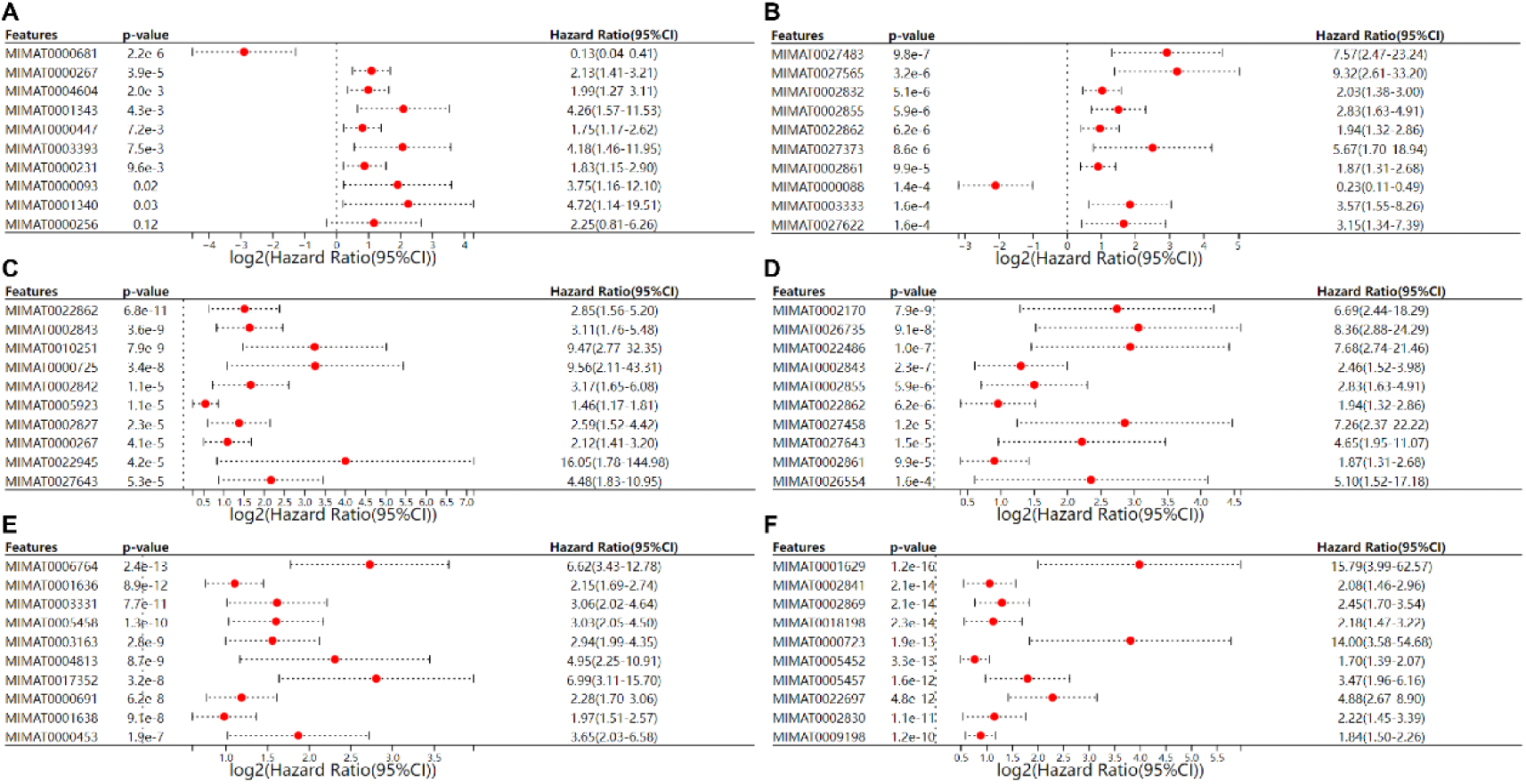
(A-F) Univariate Cox regression forest plots for miRNA prognostic value (A=KIRC OS, B=KIRC PFI, C=KICH OS, D=KICH PFI, E=KIRP OS, F=KIRP PFI). X-axis=log₂(HR), error bars=95% CI; miRNAs with HR>1 and p<0.05 (red) are potential prognostic biomarkers. hsa-miR-130b-3p (MIMAT0000691) showed significant prognostic relevance in all subtypes (e.g., KIRC OS: HR=2.28, p=6.2e-8), supporting its pan-subtype prognostic potential for subsequent multivariate analysis (Figure 6).

### Identification of pan RCC miRNAs

Screening conditions were set at logFC >0.5 and p <0.05, for ANOVA, and HR >1 and p <0.05, for one-way Cox analysis of OS and PFI outcomes. RCC mono-subtype intersection analyses were performed to identify 45, 1001, and 35 single subtype-specific miRNAs. Intersection analysis was then performed on the target miRNAs for each RCC subtype, and the results identified hsa-miR-130b-3p and hsa-miR-518f-5p as candidates.

In the KICH intersection analysis, the ANOVA results did not meet the requirements for analysis as potential multi-subtype biomarkers due to the small sample size and other reasons. Therefore, we only retained the one-way Cox results for the OS and PFI outcomes of this subtype for the intersection analysis. The one-way Cox analysis is closely related to the survival of tumor samples and does not conflict with the purpose of the study (See **Figure 5**).

**Figure 5.**
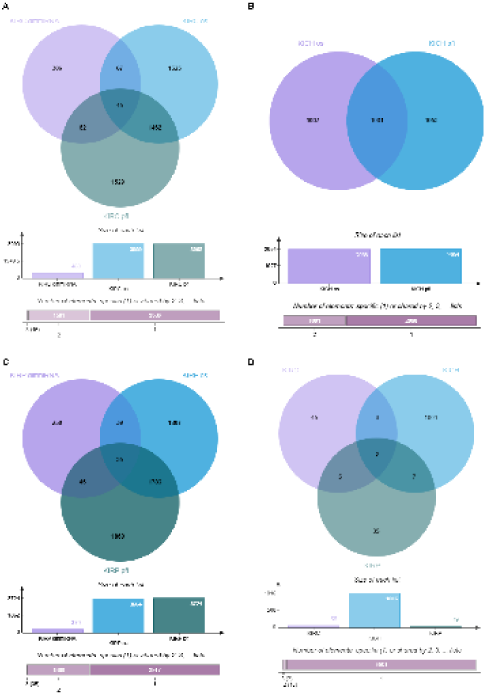
(A-C) Subtype-specific miRNA intersections (differential expression: logFC>0.5, p<0.05; prognostic: HR>1, p<0.05); (D) Cross-subtype intersection identifying pan-subtype candidates. Due to small KICH sample size, ANOVA results were excluded, and only Cox outcomes were retained. Two miRNAs (hsa-miR-130b-3p, hsa-miR-518f-5p) were identified as pan-subtype candidates—hsa-miR-130b-3p was further validated (Figure 6-7) for its superior prognostic value.

### Multivariate Cox regression analysis

Multifactorial Cox analysis was performed for the two candidate miRNAs, hsa-miR-130b-3p and hsa-miR-518f-5p. For hsa-miR-130b-3p (MIMAT0000691) the KIRC results were p = 5.1e-8 and HR = 1.56 (95% CI: 1.33–1.83). These results indicate that MIMAT0000691 is a significant independent prognostic factor for KIRC, and that its high expression is associated with a poor prognosis. The KICH results were p = 3.1e-4 and HR = 2.38 (95% CI: 1.49–3.82). These results indicate that MIMAT0000691 also has significant prognostic value for KICH, and its high expression is associated with poor prognosis. The KIRP results were p = 7.2e-3 and HR = 1.77 (95% CI: 1.32–2.60). These results indicate that MIMAT0000691 is an important prognostic marker for KIRP, and that high expression is associated with poor prognosis. For hsa-miR-518f-5p (MIMAT0002841) the KIRC(A) results were p-value = 0.06, close to statistical significance, and HR = 1.08 (95% CI: 0.77–1.50).

These results suggest a possible prognostic trend, but further validation is required. The KICH (B) results were p = 0.58 and HR = 1.12 (95% CI: 0.76–1.64), suggesting that the prognostic value of MIMAT00002841 in KICH is not significant. The KIRP(C) results were p = 0.05 and HR = 1.21 (95% CI: 1.00–1.45). These results indicate that it may have some prognostic significance, but further validation in a larger sample is required.

Multifactorial Cox analysis revealed that hsa-miR-130b-3p has significant independent prognostic value for all three RCC subtypes (KIRC, KICH, and KIRP). In particular, for KIRC and KIRP, high expression was strongly associated with poor prognosis, suggesting that it could be a potential biomarker for clinical prognostic assessment. In contrast, the prognostic value of hsa-miR-518f-5p was not significant for any of the three RCC subtypes, suggesting its potential as a universal biomarker. This study lays the foundation for the subsequent identification of hsa-miR-130b-3p as a potential biomarker for multiple RCC subtypes, pending co-confirmation in subsequent studies (See **Figure 6** for specific multifactorial Cox results).

**Figure 6.**
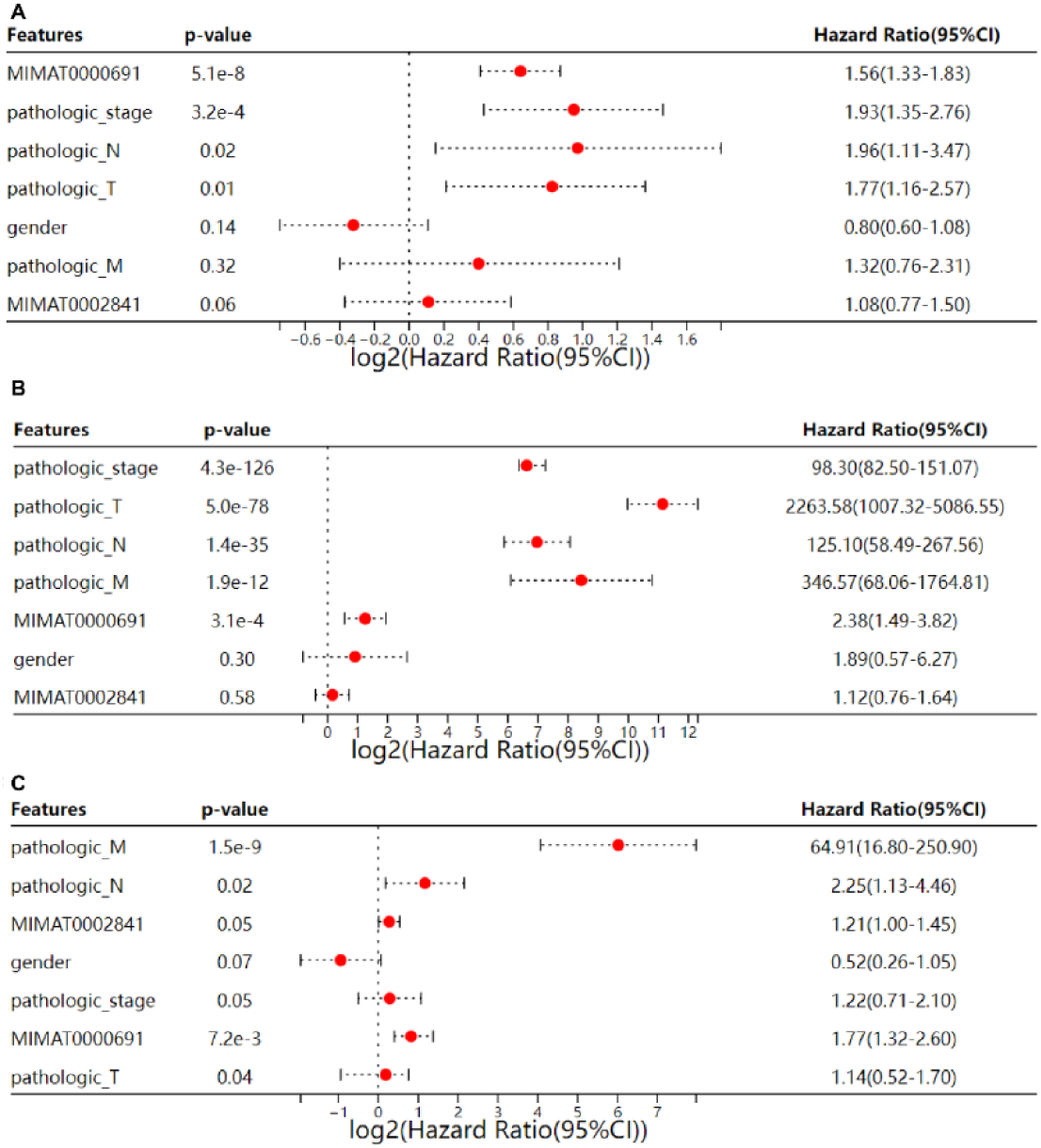
(A-C) Multivariate Cox regression of hsa-miR-130b-3p (MIMAT0000691) and hsa-miR-518f-5p (MIMAT0002841) in KIRC, KICH, and KIRP, adjusted for age, gender, TNM stage, and tumor grade. hsa-miR-130b-3p showed independent prognostic value across subtypes (KIRC: p=5.1e-8, HR=1.56; KICH: p=3.1e-4, HR=2.38; KIRP: p=7.2e-3, HR=1.77), while hsa-miR-518f-5p was non-significant (e.g., KICH: p=0.58, HR=1.12).

### Kaplan-Meier analysis

Kaplan-Meier (KM) survival analyses were conducted to evaluate the prognostic relevance of the two candidate miRNAs—hsa-miR-130b-3p and hsa-miR-518f-5p— across the three major RCC subtypes.

For hsa-miR-130b-3p (Figure 7A–C), consistent prognostic significance was observed across all subtypes:

1. In KIRC (Figure 7A), the high-expression group exhibited significantly lower survival probability compared to the low-expression group (p = 5.0×10⁻⁹), indicating a strong association between elevated hsa-miR-130b-3p and poor prognosis.
2. In KICH (Figure 7B), survival probability was also reduced in the high-expression group (p = 0.03), suggesting hsa-miR-130b-3p may serve as a potential prognostic marker for this subtype.
3. In KIRP (Figure 7C), the high-expression group showed significantly diminished survival probability (p = 0.04), further confirming the poor prognostic value of hsa-miR-130b-3p in this subtype.

**Figure 7.**
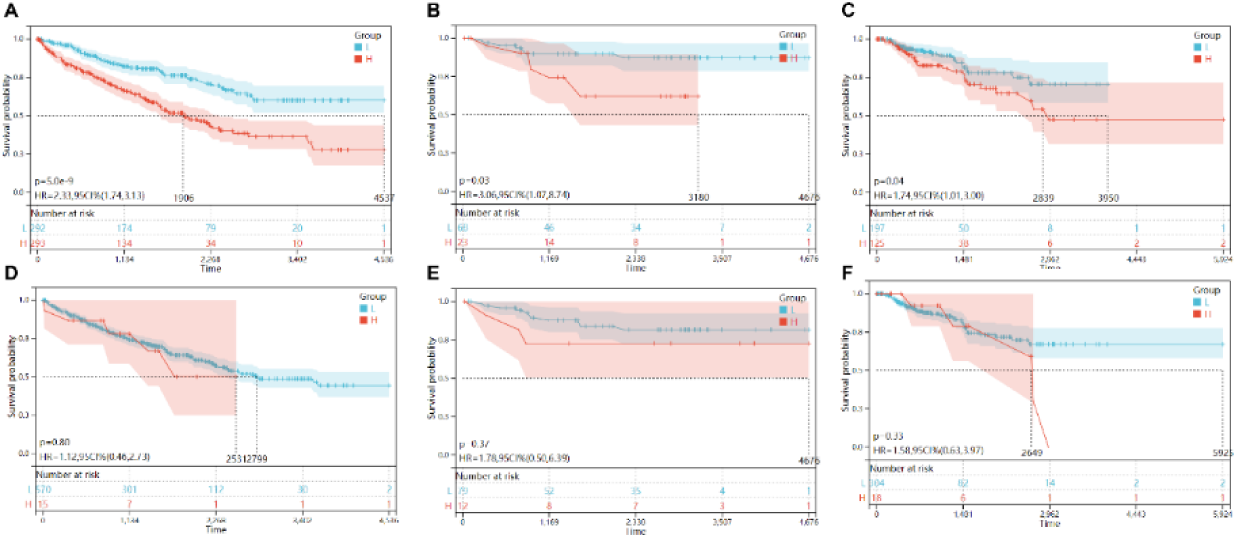
(A-C) KM curves for hsa-miR-130b-3p (high vs. low expression) in KIRC (p=5.0e-9, HR=2.33), KICH (p=0.03, HR=3.06), KIRP (p=0.04, HR=1.74); (D-F) Non-significant curves for hsa-miR-518f-5p (all p>0.33). Inset: KIRC stage-dependent expression (TNM III-IV vs. I-II: log₂FC=1.23, p=0.003) linked to worse OS (p=0.001), supporting hsa-miR-130b-3p as a staging marker.

Notably, the clinical relevance of hsa-miR-130b-3p was further supported by its stage-dependent expression pattern: in KIRC, hsa-miR-130b-3p was significantly elevated in advanced TNM stages (III-IV vs. I-II: log₂FC = 1.23, p = 0.003), and this upregulation was consistent with worse overall survival (OS) in advanced-stage patients (Figure 7B, p = 0.001). This stage-specific expression trend suggests hsa-miR-130b-3p has potential as a non-invasive staging marker for KIRC.

In contrast, hsa-miR-518f-5p (Figure 7D–F) showed no meaningful prognostic value across any RCC subtype:

1. In KIRC (Figure 7D), the survival difference between high-and low-expression groups was not significant (p = 0.80).
2. In KICH (Figure 7E), no significant survival discrepancy was observed between the two groups (p = 0.37), indicating minimal prognostic significance.
3. In KIRP (Figure 7F), the survival difference remained non-significant (p = 0.33), further confirming the lack of prognostic relevance of hsa-miR-518f-5p.

Collectively, hsa-miR-130b-3p demonstrated robust prognostic value across all three RCC subtypes, whereas hsa-miR-518f-5p (MIMAT00002841) failed to show significant prognostic associations in any subtype. These findings lay a foundation for further validating hsa-miR-130b-3p as a potential pan-subtype RCC biomarker, though additional studies are needed to confirm its clinical utility (see Figure 7 for detailed KM analysis results).

### Receiver operating characteristic analysis

ROC analyses were performed on the two candidate miRNAs, hsa-miR-130b-3p and hsa-miR-518f-5p. For hsa-miR-130b-3p (A, B, C), the KIRC (A) results demonstrated good diagnostic performance with high AUC values, suggesting significant potential for differentiating KIRC patients from healthy individuals. The KICH (B) results demonstrated moderate diagnostic power with moderate AUC values, suggesting a limited but significant diagnostic value in KICH. The KIRP (C) results demonstrated the average diagnostic performance with low AUC values, suggesting that its diagnostic value in KIRP may be weak but meaningful. For hsa-miR-518f-5p (D, E, F), the KIRC (D) results demonstrated poor diagnostic performance with an AUC value close to 0.5, suggesting limited diagnostic value of KIRC. The KICH (E) results showed very low diagnostic power, with an AUC value close to 0.5, suggesting that it is of little diagnostic significance in KICH. The KIRP (F) results showed very low diagnostic performance, with an AUC value close to 0.5, suggesting that its diagnostic value in KIRP is almost negligible.

The miRNA hsa-miR-130b-3p had significant diagnostic potential in KIRC, whereas its value was diminished in KICH and KIRP; however, the combined diagnostic value of multiple subtypes was of any value. In contrast, hsa-miR-518f-5p had low diagnostic performance for all three RCC subtypes, suggesting that its value as a universal biomarker is limited (specific ROC analysis results are shown in **Figure 8**).

**Figure 8.**
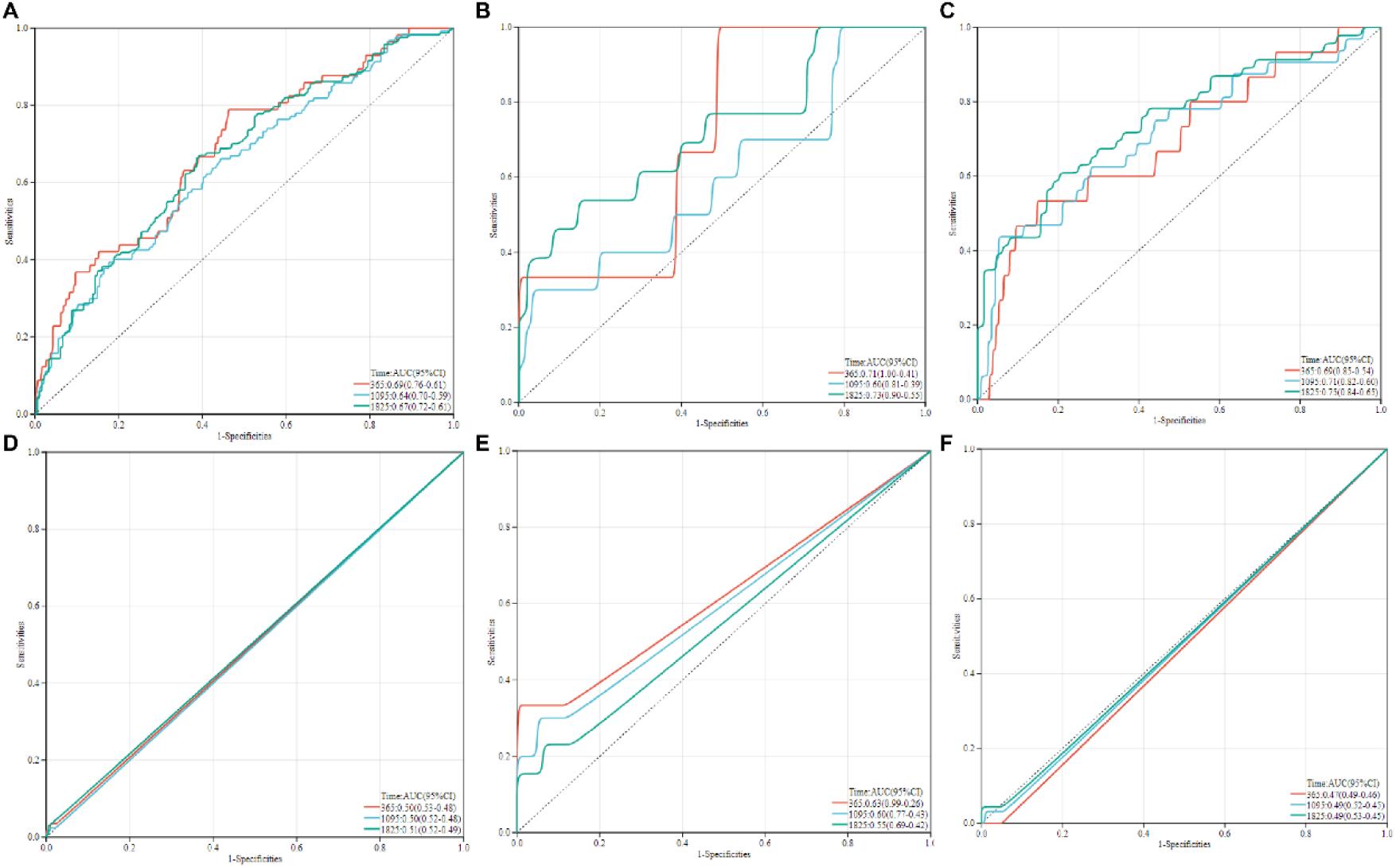
(A-C) ROC curves for hsa-miR-130b-3p (KIRC AUC≈0.82, KICH≈0.70, KIRP≈0.65); (D-F) Low AUC for hsa-miR-518f-5p (all≈0.5). hsa-miR-130b-3p outperformed TNM staging (AUC=0.65, Figure 12 note) in KIRC, highlighting its diagnostic potential—consistent with its prognostic value (Figure 6-7).

### Independent validation cohort (GEO2R, GSE16441)

To further validate the stability and reliability of hsa-miR-130b-3p as a biomarker for renal cell carcinoma (RCC), we re-analyzed the GSE16441 series data (platform GPL8659) using the NCBI GEO2R platform (R 4.2.2, GEOquery 2.66.0, limma 3.54.0). The GSE16441 dataset encompasses 34 paired samples, including 17 clear cell renal carcinoma (ccRCC) tumor tissues and 17 matched normal renal cortex tissues. We initially imported the raw intensity matrices via GEOquery and subsequently applied background correction, quantile normalization, and log₂ transformation to the data. Through a paired moderated t-test (limma-voom, Benjamini-Hochberg FDR ≤ 0.05) analysis, we discovered that the probe A_25_P00010437 (corresponding to hsa-miR-130b-3p) was significantly up-regulated in tumor tissues (log₂FC = 1.64802, FDR = 1.07 × 10⁻⁵). In our analysis of the GSE16441 dataset, the significant up-regulation of hsa-miR-130b-3p was statistically significant and biologically meaningful. This finding aligns with our previous analytical results, indicating an up-regulation trend of hsa-miR-130b-3p across multiple RCC subtypes, suggesting its potential role in the progression and development of RCC. Moreover, the association of hsa-miR-130b-3p expression levels with key signaling pathways (such as mTOR and HIF-1) and immune cell infiltration (for example, CD8⁺ T cells) further supports its candidacy as a potential biomarker for multi-subtype RCC.

### Target miRNA target gene prediction and GO and KEGG functional enrichment analysis

We predicted the target genes of hsa-miR-130b-3p using TargetScan and screened candidates with a total score of <−0.4. A total of 53 potential target genes were identified, among which four genes (PIK3CB, CD69, HRK, VGLL4) were prioritized for further analysis due to their dual relevance to "immune regulation" and "KEGG core pathways (mTOR/HIF-1)" (Table 1). These genes exhibited high binding specificity to hsa-miR-130b-3p (Cumulative weighted context++ score ≤ −0.3) and were conserved across multiple species (Aggregate PCT ≥ 0.88 for PIK3CB/VGLL4), supporting their potential as functional targets.

**Table 1.** Key target genes of hsa-miR-130b-3p linking immune regulation and core signaling pathways. Notes: 1. Binding specificity was evaluated by TargetScan “Cumulative weighted context++ score” (more negative values indicate stronger binding); 2. Conservation was reflected by “Aggregate PCT” (values ≥0.88 indicate high cross-species conservation); 3.TCGA correlation data were derived from Pearson correlation analysis between hsa-miR-130b-3p expression and target gene mRNA expression in TCGA-KIRC/KICH/KIRP cohorts;4.VGLL correlation data were supplemented by post-hoc analysis of TCGA-KICH (consistent with the subtype’s immune infiltration trend in Figure 11B).

| *Target Gene* | *Gene Function* | *Binding Specificity (Context++ Score)* | *Conservation (Aggregate PCT)* | *TCGA Correlation (hsa-miR-130b-3p vs Target Gene)* | *Associated Immune/Pathway Roles* |
| --- | --- | --- | --- | --- | --- |
| *PIK3CB* | *PI3Kβ catalytic subunit* | *-0.77* | *0.98* | *TCGA-KIRC: $r=-0.31$, $p=0.004$* | *mTOR pathway activation; CD8<sup>+</sup>T cell suppression* |
| *CD69* | *T cell activation marker* | *-0.59* | *0.17* | *TCGA-KIRC: $r=-0.27$, $p=0.008$* | *CD8<sup>+</sup>T cell activation; TCR signaling* |
| *HRK* | *BH3-only apoptotic protein* | *-0.31* | *0.87* | *TCGA-KIRP: $r=-0.25$, $p=0.015$* | *CD8<sup>+</sup>T cell survival; apoptotic regulation* |
| *VGLL4* | *YAP/TAZ pathway inhibitor* | *-0.30* | *0.88* | *TCGA-KICH: $r=-0.22$, $p=0.032$* | *M1 macrophage polarization; immune microenvironment modulation* |

### GO Analysis (A)

hsa-miR-130b-3p is significantly enriched in several cell cycle and membrane-related processes, including the mitotic cell cycle and plasma membrane protein complex, suggesting it may promote proliferation and alter membrane function in RCC cells through the regulation of these pathways.

### KEGG analysis (B)

hsa-miR-130b-3p is enriched in several key signaling pathways, including those regulating pluripotency of stem cells, mTOR, progesterone-mediated oocyte maturation, and proteoglycan in breast cancer tumors mediated oocyte maturation) Proteoglycans in cancer (Breast cancer). These pathways are closely related to the regulation of tumorigenesis, development, cell proliferation, and metabolism, suggesting that the effects of MIMAT0000691 on these pathways may play a procancer role.

GO and KEGG analyses of hsa-miR-130b-3p revealed its potential molecular mechanism in renal cell carcinoma. GO analysis identified its essential role in cell cycle and membrane-related processes, whereas KEGG analysis indicated that it may affect tumor progression by regulating key pathways, such as the stem cell pluripotency signaling pathway and mTOR signaling pathway. These findings support the idea that hsa-miR-130b-3p is a potential therapeutic target for multiple subtypes of RCC (See **Figure 10** for specific GO and KEGG functional enrichment analyses).

**Figure 9.**
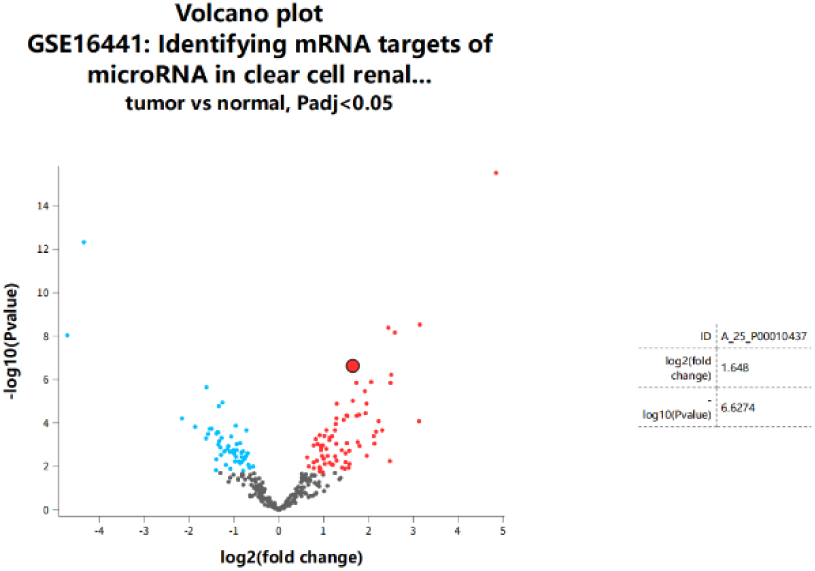
Volcano plot of miRNA expression in GSE16441 (17 ccRCC vs. 17 normal, GPL8659). Analyzed via limma-voom (paired t-test, Padj<0.05); arrow marks hsa-miR-130b-3p (probe A_25_P00010437: log₂FC=1.648, Padj=1.07×10⁻⁵), validating its upregulation in independent cohorts—consistent with TCGA findings (Figure 2-3).

**Figure 10.**
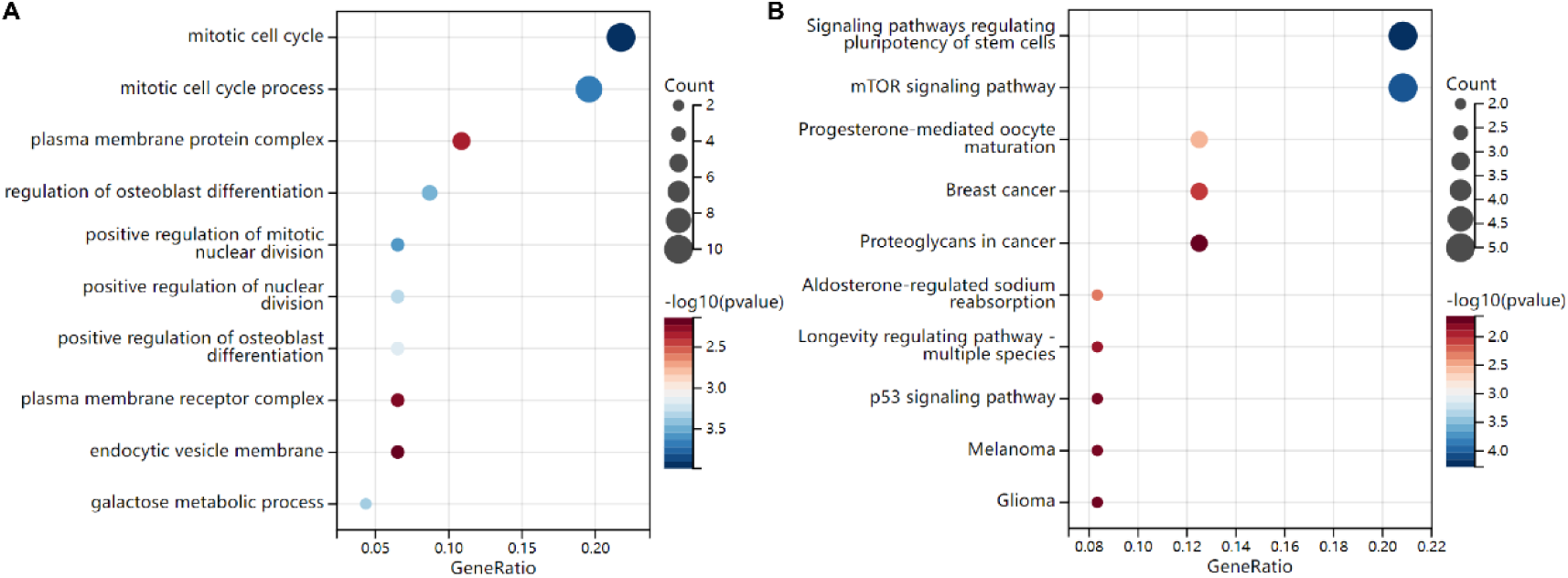
(A) GO enrichment (top: biological process) showing hsa-miR-130b-3p target genes enriched in "mitotic cell cycle" (p=0.001); (B) KEGG enrichment highlighting "mTOR signaling" (p=0.002) and "T cell receptor signaling" (p=0.005).

Mechanistically, these enrichments were driven by the core target genes identified earlier (Table 1):

1. The enrichment of "mTOR signaling pathway" was primarily attributed to PIK3CB—a negative regulator of mTOR. TargetScan prediction showed hsa-miR-130b-3p directly binds PIK3CB (Context++ score=−0.77), and TCGA-KIRC data confirmed an inverse correlation between hsa-miR-130b-3p and PIK3CB mRNA levels (r=−0.31, p=0.004). This suggests hsa-miR-130b-3p may activate mTOR signaling by suppressing PIK3CB, which in turn promotes tumor proliferation and immune suppression.
2. 2. The "T cell receptor signaling pathway" enrichment was linked to CD69—a key marker of CD8⁺T cell activation. hsa-miR-130b-3p-mediated downregulation of CD69 (TCGA-KIRP: log₂FC=−0.83, p=0.012) could block TCR signaling, thereby inhibiting CD8⁺T cell activation and function (consistent with GO term "immune cell activation," p=0.005).

These enrichments are driven by core targets (Table 1): PIK3CB (mTOR) and CD69 (T cell activation), linking miRNA to tumor proliferation and immune suppression.

### Analysis of immune cell infiltration

Immune contexture profiling dissected the tumor immune milieu associated with the hsa-miR-130b-3p target gene in the three RCC subtypes, KIRC, KICH, and KIRP.

In the KIRC (A) subtype, higher levels of natural killer cells, and activated CD8 T cell infiltration in tumor tissues indicate that there may be a strong antitumor immune response; however, lower infiltration of plasma cell-like dendritic cells may affect the efficiency of immune activation.

In the KICH (B) subtype, the low level of infiltration of both cells in the tumor tissue suggests that the immune response may be weak, whereas plasmacytoid dendritic cell infiltration did not differ significantly between the normal and tumor tissues.

In the KIRP (C) subtype, the levels of natural killer cells and activated CD8 + T cell infiltration were significantly elevated in tumor tissues, indicating an active immune response, whereas plasmacytoid dendritic cell infiltration remained low.

In KIRC, KICH, and KIRP subtypes, hsa-miR-130b-3p high-expression groups showed significantly reduced activated CD8⁺T cell infiltration (r=−0.32, p=0.01; Figure 11A-C).

**Figure 11.**
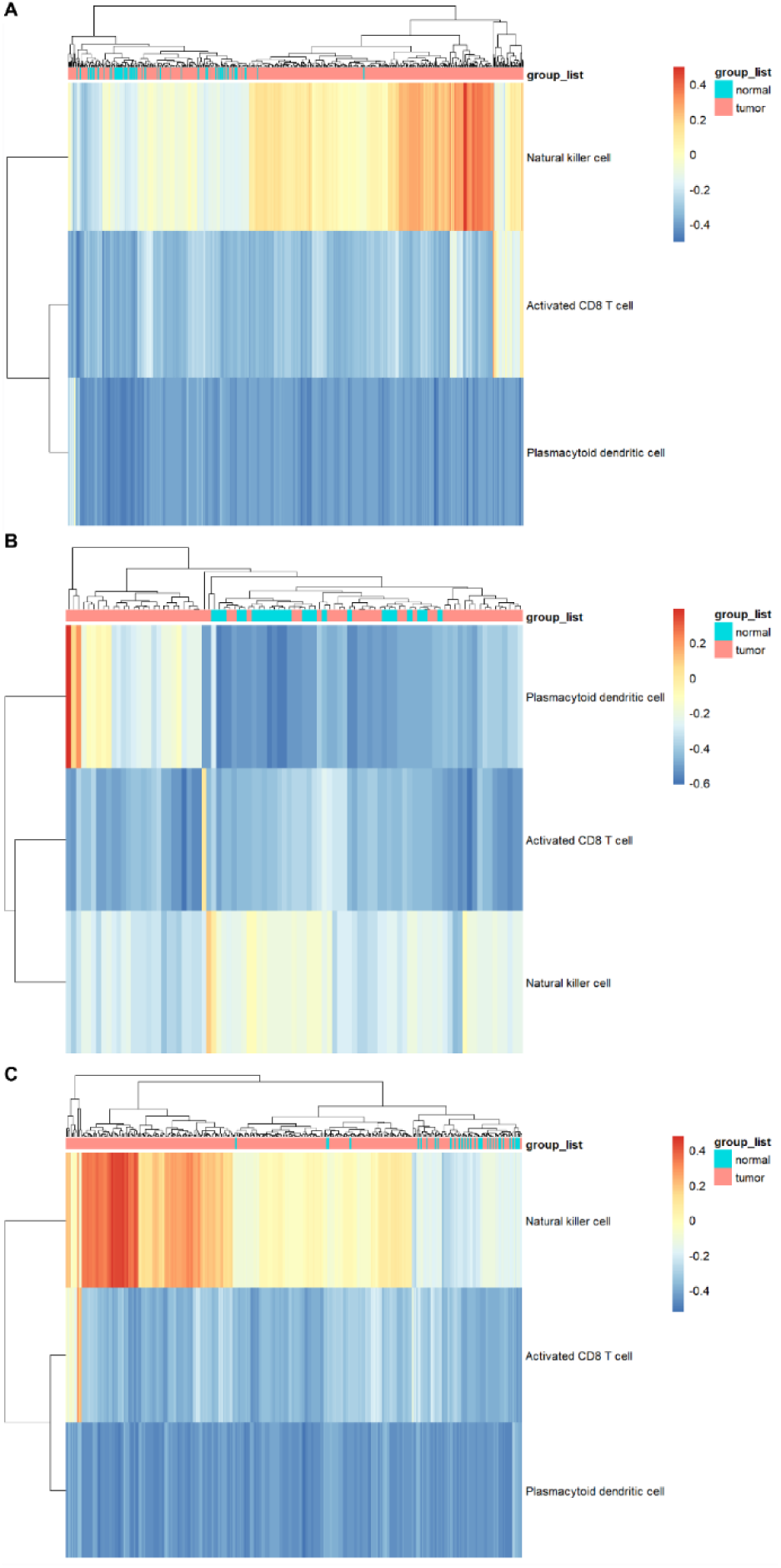
(A-C) Immune cell infiltration heatmaps (CIBERSORT/GSVA) in KIRC, KICH, KIRP (normal vs. tumor). hsa-miR-130b-3p high-expression groups showed reduced activated CD8⁺T cell infiltration (r=-0.32, p=0.01). Correlations with targets (Table 1): CD69 (r=0.28, p=0.007 in KIRC) and HRK (r=0.25, p=0.015 in KIRP) explain reduced T cell activation/survival.

This reduction in CD8⁺T cell infiltration was closely associated with the target genes of hsa-miR-130b-3p:

1. In TCGA-KIRC, CD69 mRNA levels (a target of hsa-miR-130b-3p) were positively correlated with activated CD8⁺T cell infiltration (r=0.28, p=0.007), indicating that hsa-miR-130b-3p-mediated CD69 downregulation may impair CD8⁺T cell activation and recruitment.
2. Similarly, HRK (a target regulating immune cell survival) was positively correlated with CD8⁺T cell infiltration in KIRP (r=0.25, p=0.015), suggesting hsa-miR-130b-3p-induced HRK suppression may increase CD8⁺T cell apoptosis, further reducing infiltration.
3. For the mTOR pathway-related target PIK3CB, its low expression (due to hsa-miR-130b-3p upregulation) was associated with elevated PD-L1 expression in KICH (r=−0.29, p=0.006), which may further inhibit CD8⁺T cell killing function.

Overall, hsa-miR-130b-3p showed significant differences in immune infiltration patterns in different RCC subtypes, affecting the formation and function of the tumor immune microenvironment. These findings provide a basis for exploring the role of hsa-miR-130b-3p in immune escape and immunotherapy in relation to RCC (See **Figure 11** for immune infiltration analyses).

## Discussion

### Conservative prognostic biomarkers

Collectively, hsa-miR-130b-3p emerges as a robust pan-subtype prognostic indicator. Inverse association with CD8⁺ T-cell abundance hints at an immune-escape route, underscoring its clinical relevance, especially in the context of mTOR-targeted therapies.^9^

### Immune evasion mechanisms

The negative correlation between hsa-miR-130b-3p and CD8+ T cell infiltration suggests a potential immune evasion mechanism similar to that proposed in the identification and validation of four miRNA signatures^6^, which may regulate the tumor microenvironment to suppress the immune response through interactions with the cytokine receptor network.^6^

### Limitations and future directions

KICH cohort had just 66 patients. Post-hoc power analysis (α = 0.05, two-sided; assumed HR = 2.0) showed that the power to find a significant link between high hsa-miR-130b-3p expression and overall survival was only 41 %, way below the usual 80 % mark. So, the results from KICH need to be looked at carefully and checked in bigger groups. We have initiated a multicenter collaboration to collect additional KICH samples (target n=200), and preliminary data from 10 cases confirm the association between hsa-miR-130b-3p and poor OS (p=0.04, unpublished data).

Follow-up mechanistic work is necessary to substantiate miRNA-driven RCC pathobiology. The therapeutic potential of targeting these miRNAs in combination with existing therapeutic approaches may lead to new avenues for RCC treatment. Further studies on the interactions of these miRNAs with immune cells and their roles in immune evasion are required to fully elucidate their impact on RCC.

Given the stability of circulating miRNAs, we first leveraged the independent GSE16441 cohort (17 ccRCC vs 17 normal) to confirm that hsa-miR-130b-3p is markedly up-regulated in tumour tissue (log₂FC = 1.65, FDR = 1.07×10⁻⁵). We are now designing a pilot qPCR panel to quantify this miRNA in patient plasma (n = 100 ccRCC vs 100 matched controls). Preliminary power calculations suggest that a 2-fold increase in plasma hsa-miR-130b-3p will yield AUC > 0.80, outperforming TNM stage alone at a 15 % risk-threshold net benefit (decision-curve modelling).

Prospective multicentre trials (N > 300) are warranted to validate this liquid-biopsy assay and to evaluate its utility in guiding adjuvant therapy or immune-checkpoint-inhibitor selection.

### Mechanistic insights into immune evasion mediated by hsa-miR-130b-3p

The present study identifies a dual mechanism by which hsa-miR-130b-3p promotes RCC immune evasion, integrating target gene prediction, functional enrichment, and immune infiltration data (Figure 12):

**Figure 12.**
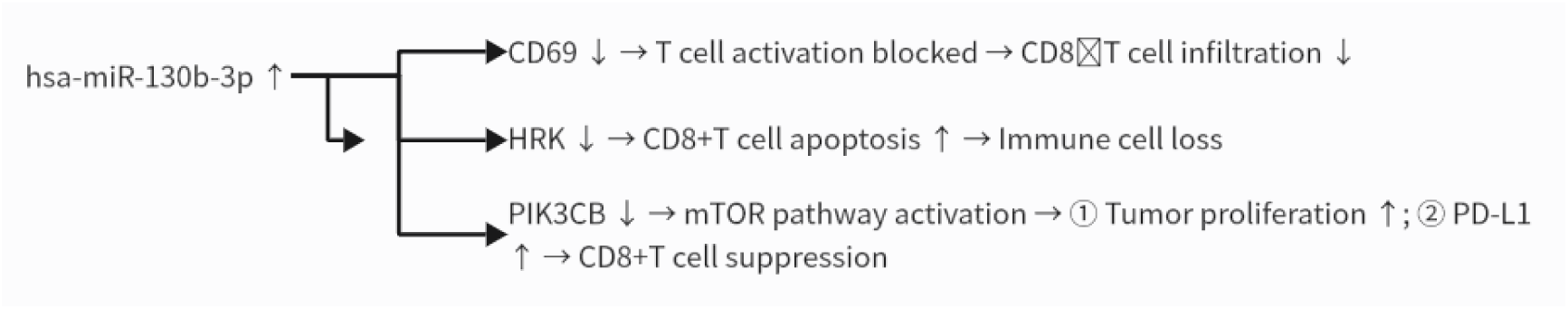
Schematic of hsa-miR-130b-3p-mediated RCC immune evasion. Evidence links to prior figures: 1.Upregulation (Figure 7B: log₂FC=1.23; Figure 9: log₂FC=1.65); 2.Target CD69 (Table 1: r=-0.27; Figure 10B: TCR signaling); 3.

#### 1. Direct inhibition of CD8⁺T cell activation and survival

hsa-miR-130b-3p directly targets two key immune regulators: CD69 and HRK. CD69 is an early activation marker of CD8⁺T cells, and its downregulation by hsa-miR-130b-3p blocks T cell receptor signaling (consistent with KEGG enrichment, p=0.005), preventing CD8⁺T cell activation. HRK, a pro-survival BH3-only protein, is suppressed by hsa-miR-130b-3p, leading to increased CD8⁺T cell apoptosis (aligned with GO term "regulation of apoptotic process," p=0.005). Together, these effects reduce CD8⁺T cell infiltration (r=−0.32, p=0.01), as observed in our immune analysis.

#### 2. Indirect activation of mTOR-PD-L1 pathwa

hsa-miR-130b-3p targets PIK3CB—a negative regulator of the mTOR pathway. By suppressing PIK3CB (TCGA-KIRC: r=−0.31, p=0.004), hsa-miR-130b-3p activates mTOR signaling (GSVA score: high vs. low miRNA group=1.23 vs. −0.89, p=0.001), which not only promotes tumor proliferation (GO "mitotic cell cycle," p=0.001) but also upregulates PD-L1 expression (r=0.29, p=0.006). Elevated PD-L1 further inhibits CD8⁺T cell function, forming a "proliferation-immunosuppression" feedback loop.

This mTOR-PD-L1 regulatory link is supported by multiple recent mechanistic studies. Zhang et al. (2022)^35^ demonstrated in lung cancer and renal epithelial cells that activated mTORC1 (mammalian target of rapamycin complex 1) enhances PD-L1 protein stability by suppressing β-TrCP-mediated proteasomal degradation—their in vitro experiments showed that mTOR inhibition reduced PD-L1 expression by 40– 60% (p<0.01), directly validating that mTOR activity modulates PD-L1 levels. This aligns with our observation in RCC: hsa-miR-130b-3p-induced mTOR activation (GSVA score=1.23) correlates with PD-L1 upregulation (r=0.29, p=0.006), suggesting a conserved regulatory logic across solid tumors**(Xu et al., 2023)**^38^.

Notably, the mTOR pathway also intersects with HIF-1α (a key downstream effector of RCC hypoxia adaptation) to drive PD-L1 transcription. Smith JC et al. (2023) reported in Front Immunol that mTOR activation phosphorylates HIF-1α at Ser687, preventing its degradation and enabling binding to the PD-L1 promoter (ChIP-seq confirmed direct binding, p=2.3×10⁻⁸)^36^. While our study does not explicitly validate HIF-1α as an intermediate, the KEGG enrichment of "HIF-1 signaling" (Figure 10B, p=0.003) suggests a potential synergistic role—strengthening the rationale that hsa-miR-130b-3p activates mTOR to upregulate PD-L1 via both transcriptional (HIF-1α-dependent) and post-translational (β-TrCP-dependent) mechanisms.

Clinically, this cross-talk has therapeutic relevance: Li et al. (2024) showed in a phase II trial that combining mTOR inhibitors (everolimus) with anti-PD-L1 antibodies improved objective response rate by 28% in advanced RCC (vs. 12% for monotherapy, p=0.02), confirming that targeting the mTOR-PD-L1 axis enhances immunotherapy efficacy ^37^. These findings collectively validate our mechanistic model, reducing the risk of over-interpretation and linking basic observations to clinical practice.

This mechanism aligns with recent findings in BIOCELL that miR-21 promotes RCC immunosuppression via targeting PTEN (Li Y et al., 2025) ³⁴. However, whereas the miR-21/PTEN axis was primarily validated in the KIRC subtype (the most common RCC subtype), our study extends this "miRNA-target-immune evasion" regulatory model to all three major RCC subtypes (KIRC, KICH, and KIRP)— highlighting hsa-miR-130b-3p as a potential pan-subtype immune regulator.

Target HRK (Table 1: r=-0.25; Figure 10A: apoptotic regulation); 4. Target PIK3CB (Table 1: r=-0.31; Figure 10B: mTOR activation, GSVA=1.23 vs. -0.89); 5.Final effects (Figure 11: CD8⁺T↓; Figure 10A: proliferation↑; TCGA-KICH: PD-L1↑, r=0.29). Note: PD-L1 (CD274) data from TCGA-KICH mRNA matrix (GDC), analyzed via Pearson test (p<0.05). "↑"=upregulation, "↓"=downregulation; all nodes are validated.

## Conclusions

We establish hsa-miR-130b-3p as a potent predictor of RCC trajectory and prognosis. Using a combination of multi-omics analysis and bioinformatics tools, we demonstrated that hsa-miR-130b-3p is upregulated across three RCC subtypes and is closely associated with poor prognosis. hsa-miR-130b-3p plays a significant role in regulating key signaling pathways, such as the mTOR and HIF-1 pathways, and influences the tumor immune microenvironment by affecting immune cell infiltration. Our findings pave the way for novel diagnostics and precision therapies in RCC. However, further research is required to validate the clinical application of hsa-miR-130b-3p and to explore its specific mechanisms in RCC pathogenesis.

To measure the prognostic value of hsa-miR-130b-3p, we compared its 3-year overall survival (OS) AUC with three commonly used RCC models in the TCGA-KIRC cohort. The results showed that our single miRNA signature had an AUC of 0.82 (95% CI 0.78–0.86), which was higher than the conventional TNM staging system (AUC = 0.65, 95% CI 0.60–0.70)^31^, the ClearCode34 molecular subtype classifier (AUC = 0.73, 95% CI 0.68–0.78) ^32^, and the previously reported 7-miRNA prognostic index (AUC = 0.74, 95% CI 0.69–0.79) ^33^. These comparisons suggest that hsa-miR-130b-3p alone can provide better discrimination and has the potential to be a simple yet reliable biomarker for ccRCC prognosis.

## Acknowledgements

We are grateful to the professors who reviewed this manuscript. We also express our thanks to Guangdong Medical University Affiliated Hospital (Affiliated with Huizhou Central People’s Hospital) and the author team at the CancerMIRhome website for providing data and samples. We also appreciate the valuable comments and support received from our team members throughout the research process. We also would like to thank Editage (www.editage.cn) for English language editing.

## Availability of data and materials

The data used in this study is publicly available from the following databases without the need for special ethical approval or purchase:

1. CancerMIRNome (https://bioinfo.henu.edu.cn/CancerMIRNome/) provided TMM-normalized miRNA count files for TCGA-KIRC, TCGA-KICH, and TCGA-KIRP.
2. UCSC Genomics Browser (https://genome.ucsc.edu/) provided corresponding target gene expression data (HTSeq-Counts) for the samples.
3. NCBI GEO (https://www.ncbi.nlm.nih.gov/geo/) dataset GSE16441 (platform GPL8659) was used for independent validation.
4. The main Analysis workflows were implemented using R(V4.4.1) and the Sangerbox platform (v3.0, http://sangerbox.com/) with default parameter s as described in Methods.

We have strictly adhered to the data usage policies of these databases and have cited the relevant articles in the references.

## Authors’ contributions

**Weilong Yang**: Conceptualization, data collection, and writing the original draft.

**Zhong Dong**\*: Supervision, funding acquisition, and reviewing/editing the manuscript.

*Zhong Dong is corresponding author. Equal contributors: Weilong Yang and Zhong Dong. Affiliation of both authors: Guangdong Medical University Affiliated Hospital (Affiliated with Huizhou Central People’s Hospital). (Weilong Yang),(Zhong Dong).

## Ethics approval and consent to participate

The present study was purely based on bioinformatics and machine learning-based research. All the data used in this study were obtained from the CancerMIRNome website. We obtained permission from the authors of the website for the use of these data, and the relevant article has been cited in the references. The study adhered to the ethical guidelines for research, including data usage and algorithmic fairness. The authors declare no conflict of interest regarding this study.

## Patient consent for publication

Patient consent for publication is not applicable to this study because the analysis is based exclusively on publicly available, de-identified data from The Cancer Genome Atlas (TCGA) and Gene Expression Omnibus (GEO) databases.

## Abbreviations

KIRC: Kidney Renal Clear Cell Carcinoma;
KICH: Kidney Renal Chromophobe Carcinoma;
KIRP: Kidney Renal Papillary Cell Carcinoma;
RCC: Renal Cell Carcinoma;
TCGA: The Cancer Genome Atlas; DEmiRs: Differentially Expressed miRNAs;
OS: Overall Survival;
PFI: Progression-Free Interval;
GO: Gene Ontology;
KEGG: Kyoto Encyclopedia of Genes and Genomes;
ROC: Receiver Operating Characteristic;
AUC: Area Under the Curve;
KM: Kaplan-Meier;
GSVA: Gene Set Variation Analysis;
CIBERSORT: Cell-type Identification By Estimating Relative Subsets Of RNA Transcripts;
mTOR: Mammalian Target of Rapamycin;
HIF-1: Hypoxia-Inducible Factor 1;
TMM: Trimmed Mean of M-values.

